# Energetic profiling reveals thermodynamic principles underlying amyloid fibril maturation

**DOI:** 10.1101/2025.05.14.653959

**Authors:** Ramon Duran-Romaña, Joost Schymkowitz, Frederic Rousseau, Nikolaos Louros

## Abstract

Amyloid fibrils adopt diverse structural polymorphs that underlie disease-specific phenotypes, but the thermodynamic principles guiding their formation and maturation remain poorly understood. Here, we apply energetic profiling to structural time series from cryo-EM datasets of IAPP, tau, and α-synuclein to decode the principles governing fibril maturation, stability and polymorphic divergence. By mapping residue-level free energy contributions across experimentally resolved assembly pathways, we reconstruct the complex maturation trajectories of distinct amyloid systems. We find that amyloid assembly is driven by aggregation-prone regions that act as sequence-encoded stabilizing motifs. As assembly progresses, these motifs are reorganized and expanded, while additional regions introduce structural frustration that enables conformational flexibility. In several cases, environmental cofactors such as metal ions or polyanions are observed in association with regions of structural remodeling, where they may act to compensate for otherwise energetically strained conformations. Notably, structurally distinct polymorphs can exhibit similar energetic profiles, while polymorphs with similar folds may differ thermodynamically depending on their assembly environment. This framework connects polymorph structure to both encoded sequence features and extrinsic modifiers, offering mechanistic insight into how amyloid strains mature and diversify over time.

## Introduction

Amyloid fibrils are a common form of protein aggregation^1^, linked to over fifty human diseases, including Alzheimer’s, Parkinson’s, and type 2 diabetes^2^. Despite arising from proteins with widely different sequences and biological roles, these aggregates share a hallmark structural feature: the cross-β architecture^3^. In this configuration, protein monomers form β-strands that stack into extended sheets, stabilized by backbone hydrogen bonds along the fibril axis. These sheets are further packed together through tightly interlocking side chains, forming so-called steric zippers that define the protofilament core and mediate interactions between protofilaments^4^. Notably, many amyloid fibrils are stabilized by short, hydrophobic sequence motifs known as aggregation-prone regions (APRs), which provide nucleation sites for fibril growth and often form the energetic spine of the fibril core^5-8^.

In addition to structural stability, the cross-β structure also enables amyloid fibrils to propagate in a prion-like manner. In this process, existing fibrils act as templates that induce the misfolding and incorporation of soluble monomers. This seeding mechanism supports sustained fibril growth, transmission between cells, and, in some cases, even spread between organisms^9-12^. Crucially, it also allows fibrils to replicate their specific conformations, giving rise to distinct structural strains, or polymorphs, with potentially different biological effects^13,14^.

Recent advances in high-resolution cryo-electron microscopy (cryo-EM) have provided atomic-level insight into these structures, revealing near-atomic models for hundreds of amyloid fibrils and uncovering an unexpected diversity of structural polymorphs.^15-17^. Strikingly, a single protein can adopt multiple discrete fibril folds that correlate with specific clinical phenotypes. For example, structurally distinct tau fibrils have been isolated from brains of patients with different tauopathies, yet each disease consistently exhibits a conserved fold across individuals^18^. Similar correlations have been observed for amyloid-β^19-21^, α-synuclein^22,23^, TDP-43^24,25^, and other amyloid-forming proteins^26,27^. However, fibrils formed in vitro frequently fail to replicate the structures found in patient tissue^17,28,29^, pointing to important, but still poorly understood, roles for cellular cofactors^29^, interactions with other biomolecules^16,30^, and post-translational modifications in shaping fibril structure^31^.

This growing structural diversity has raised new questions about how sequence and environment determine amyloid fibril structure, and how these polymorphs form and evolve over time^6,32-34^. To address this, we recently developed an in silico method to calculate residue-level energetic stability of fibril structure^6^, enabling systematic classification of disease-associated polymorphs based on their energetic profile^33^. This method was later independently validated by its ability to predict experimental measures of fibril stability and assembly^35^. Building on this method, we now apply energetic profiling to published structural time series of amyloid fibrils formed by IAPP^36^, tau^37^, and α-synuclein^38^. Remarkably, our approach reconstructs the complex maturation timelines of each protein, uncovering the energy landscape that governs their folding, assembly, and polymorphic divergence. We find that fibril assembly is anchored by APRs that act as sequence-encoded stabilizing motifs. As assembly progresses, these motifs undergo reorganization, while other segments introduce structural frustration that enables conformational diversification. In several cases, environmental cofactors, such as metal ions or polyanions, are found in conjunction with structural remodeling, where they appear to compensate for regions of local energetic strain and enable otherwise unfavorable conformations. Together, these findings reveal a unifying thermodynamic framework in which intrinsic sequence features and extrinsic factors collectively shape the structural diversity of amyloid strains observed in disease.

## Results

### APR stabilization underlie the emergence of distinct IAPP fibril polymorphs

Time-resolved cryo-EM of fibrils assembled in vitro from the human islet amyloid polypeptide (IAPP) mutant S20G recently revealed notable structural diversity during assembly^36^. Early-stage fibrils begin as a shared protofilament with a P-shaped cross-section, which later branches into two distinct lineages: one forming C-shaped and the other L-shaped protofilament cores (Fig. 1A). To understand the forces underlying their structural maturation, energetic profiling reveals that the late-stage fibril cores of both lineages become significantly more stable than their shared P-shaped intermediate (Fig. 1B, C), nearly doubling the average per-residue stabilizing free energy, while the added peripheral protofilaments remain comparatively unstable. This increased stability of the fibril cores is not uniformly distributed. Instead it is concentrated within three APRs previously identified as critical for IAPP aggregation, spanning residues 12-17^39,40^, 22-27^41,42^, and 30-36^43,44^, and located within β-strand regions that maximize backbone hydrogen bonding (Fig. 2A and Fig. 1D). As fibrils mature, these APRs not only become more stabilizing but also expand their structural contribution to the fibril core, a process facilitated by the formation of additional hydrogen bonds that reinforce β-sheet stacking (Fig. 1D, E and Fig. S1). This is further reinforced by strengthened side-chain packing, enhanced hydrophobic burial, and more favorable solvation (Fig. 1F-H). In contrast, the sequences between APRs remain relatively unstable or become more frustrated, contributing little or negatively to the fibril’s overall stability (Fig. 2A). This produces a biphasic energy landscape composed of localized stabilizing hotspots embedded within frustrated or neutral regions (Fig. 1C). Consistent with thermodynamic expectations, these patterns suggest that fibril maturation proceeds along an energetically downhill trajectory, with increasing stabilization concentrated in key APR motifs as polymorphs evolve toward distinct yet thermodynamically favorable endpoints.

**Figure 1.**
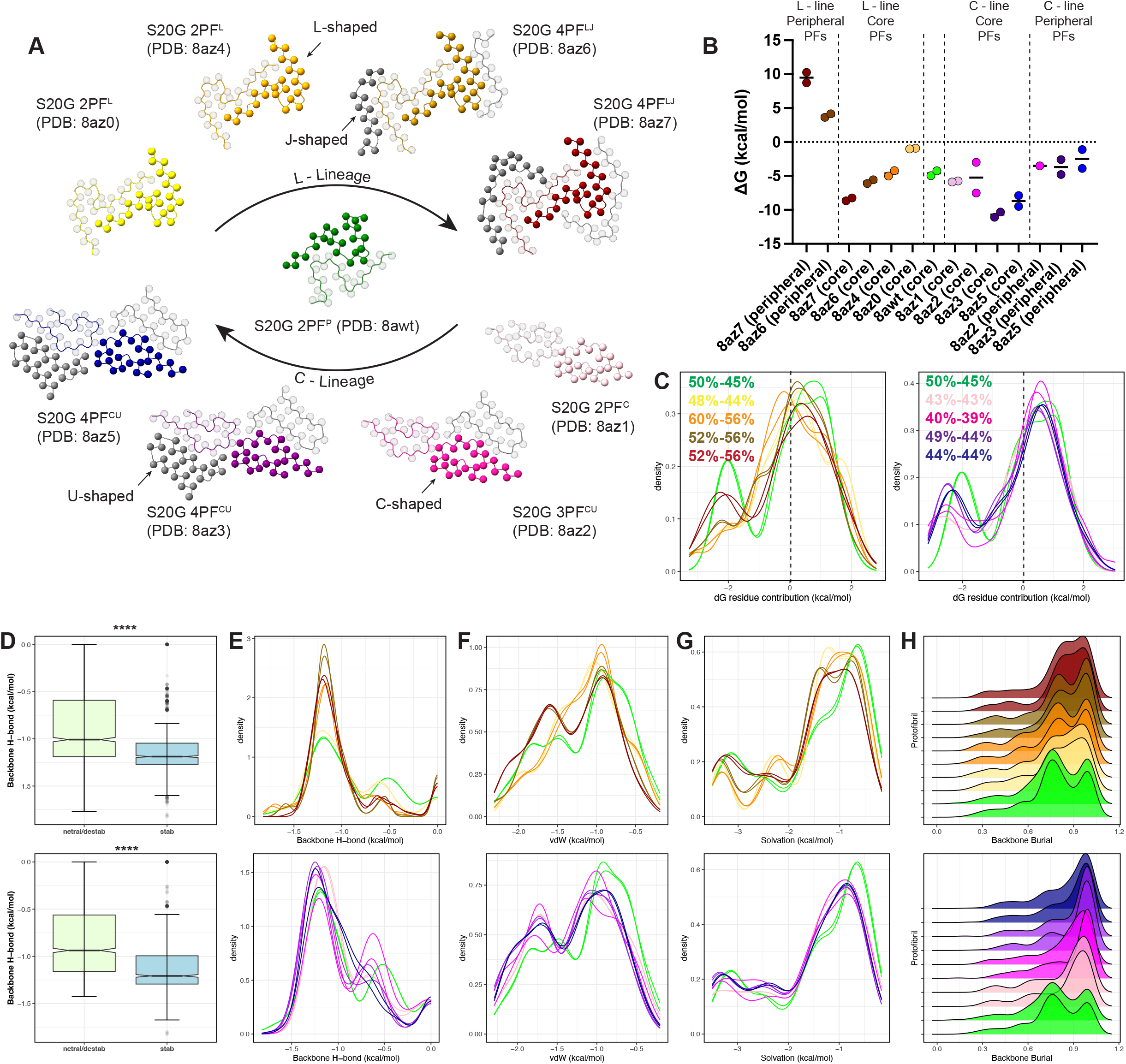
Structural assessment of IAPP intermediate protofilaments. (A) Graphical representation of amyloid fibrils formed by the S20G IAPP mutant. Following the formation of an early intermediate (shown in green), two separate structural lines are generated as kinetics of aggregation progress (L- and C-lineage). In each structure, colored protofilaments represent the core structural elements that define the fibril polymorph, while grey protofilaments denote later-added peripheral elements that contribute less to the overall stability (B) Total stability measurements of protofilament structures. (C) Distribution of individual residue energy contributions to protofilament stabilities. Percentage of residues contributing to protofilament stability are shown. (D) Backbone hydrogen-bond energies of stabilizing (shown in cyan) and neutral or destabilizing residues (shown in light green). Statistics: t-test comparison of means. (E-H) Distribution of per residue contributions to (E) backbone H-bonds, (F) van der Waals interactions, (G) solvation energies, as well as (H) total side-chain burial values. All plots are split per lineage and color-coded as in A.

**Figure 2.**
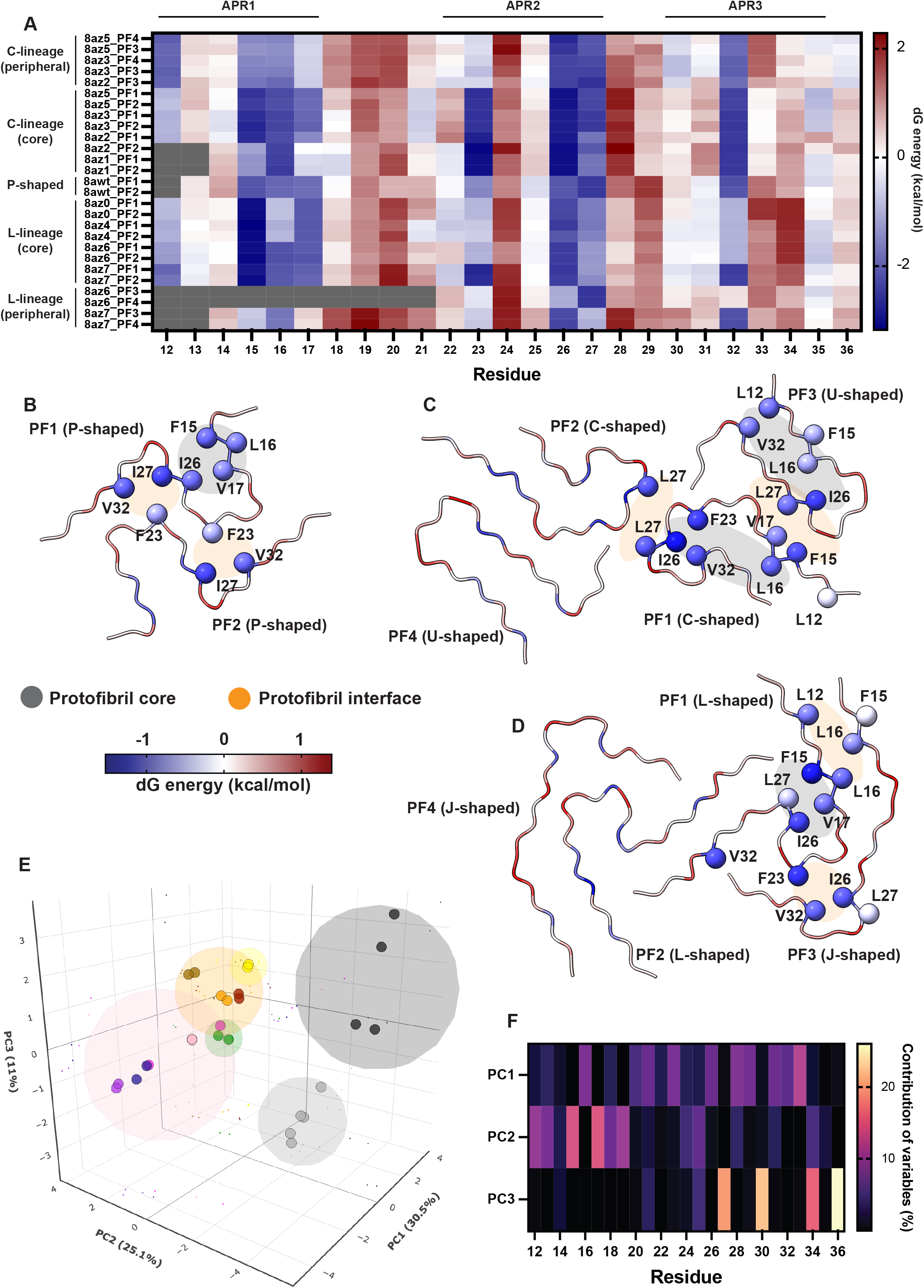
Thermodynamic profiling of IAPP S20G intermediate protofilaments. (A) Individual residue contributions (residue numbers shown in x-axis) to the stability of protofilaments. Protofilament profiles are clustered based on evolution of both lineages (shown left). Experimentally determined APRs are highlighted at the top. (B-D) Different favorable interactions between common residues stabilize the different protofilament cores (highlighted in grey) and cross-protofilament interfaces (highlighted in orange). (E) Principal component analysis and k-means clustering of IAPP protofilament thermodynamic profiles reconstructs the C-lineage (pink cluster) and L-lineage (orange cluster). The common early P-shaped intermediate is located between the two clusters in the eigenspace (green cluster), with peripheral fibrils forming separate distant clusters (grey-shaded clusters). Protofilaments are color-coded as in Figure 1A. (F) Per residue contributions to the main principal components.

Despite their morphological divergence, the mature C- and L-folds are anchored by the same aggregation-prone regions, but each lineage reinforces different parts of the sequence (Fig. 2A). The L-fold enhances stability in the N-terminal APR, while the C-fold draws more stabilizing energy from the central APR, consistent with its known role in IAPP nucleation. These findings indicate that sequence-encoded stabilizing motifs can be differentially deployed across lineages, producing distinct structural outcomes that are thermodynamically comparable but stabilized through alternative energetic routes. Further structural analysis revealed that a set of eight hydrophobic residues (L12, F15, V17, F23, I26, L27, V32, and F36) recurs across all folds and plays a central role in core stabilization. In the P- and L-folds, a subset of these residues bridge the N-terminal and central APRs through steric zippers packed within the protofilament core, with the rest stabilizing protofilament contacts. In the C-fold, however, the same residues shift to stabilize the core and inter-protofilament contacts (Fig. 2B–D). This flexible reuse highlights how a single sequence can support multiple stable folds by adapting its packing interfaces to different structural contexts.

To assess global energetic trends, principal component analysis (PCA) of residue-level energy profiles confirms these relationships. The intermediate P-shaped fold occupies a position between the mature C and L clusters in the energy space, while more peripheral protofilaments form outliers (Fig. 2E). The first PCA component, driven by changes in the central APR, clearly separates the two lineages. Analysis of variance confirms that APRs account for most of the energetic differentiation observed during fibril evolution (Fig. 2F).

### Backbone strain accumulates during tau fibril maturation despite APR-driven stabilization

We next asked whether the patterns observed for IAPP fibril maturation extend to other amyloid systems. Tau fibrils assembled in vitro from residues 293–391 can adopt disease-relevant folds resembling those found in Alzheimer’s disease (AD) and chronic traumatic encephalopathy (CTE). Remarkably, both polymorphs can be reproduced under simplified conditions by varying only the buffer composition, specifically using Mg^2+^ to promote AD-like folds and NaCl to favor CTE-like structures^37,45^. This minimal system provides a rare opportunity to dissect how intrinsic thermodynamic forces govern amyloid maturation, independent of cellular cofactors or post-translational modifications. Recent time-resolved cryo-EM reconstructions of tau protofilaments assembled under these conditions offer direct insight into the conformational transitions and energetic constraints that shape their maturation pathways (Fig. S2).

Energetic profiling of the tau time-series reveals a similar biphasic pattern (Fig. 3A, B, Fig. S3B and Fig. S4B). As assembly progresses, however, tau fibrils show a progressive loss of overall stability that contrasts with the IAPP behavior: the number of destabilizing residues steadily increases across both AD and CTE lineages (Fig. 3C, D). This thermodynamic penalty originates almost exclusively from backbone hydrogen bonds that become increasingly strained (Fig. 3E, F, Fig. S3A and Fig. S4A), leading to a progressive loss of stabilizing energy contributions. As a result, we observe a strong negative correlation between fibril maturation stage and the magnitude of backbone hydrogen-bond energy (Fig. 3G, H). Other interactions, such as hydrophobic burial and solvation, remain largely unchanged (Fig. S3D, E and Fig. S4D, E), indicating that the expanding tau fibril cores tighten at the cost of local backbone geometry rather than through loss of side-chain packing.

**Figure 3.**
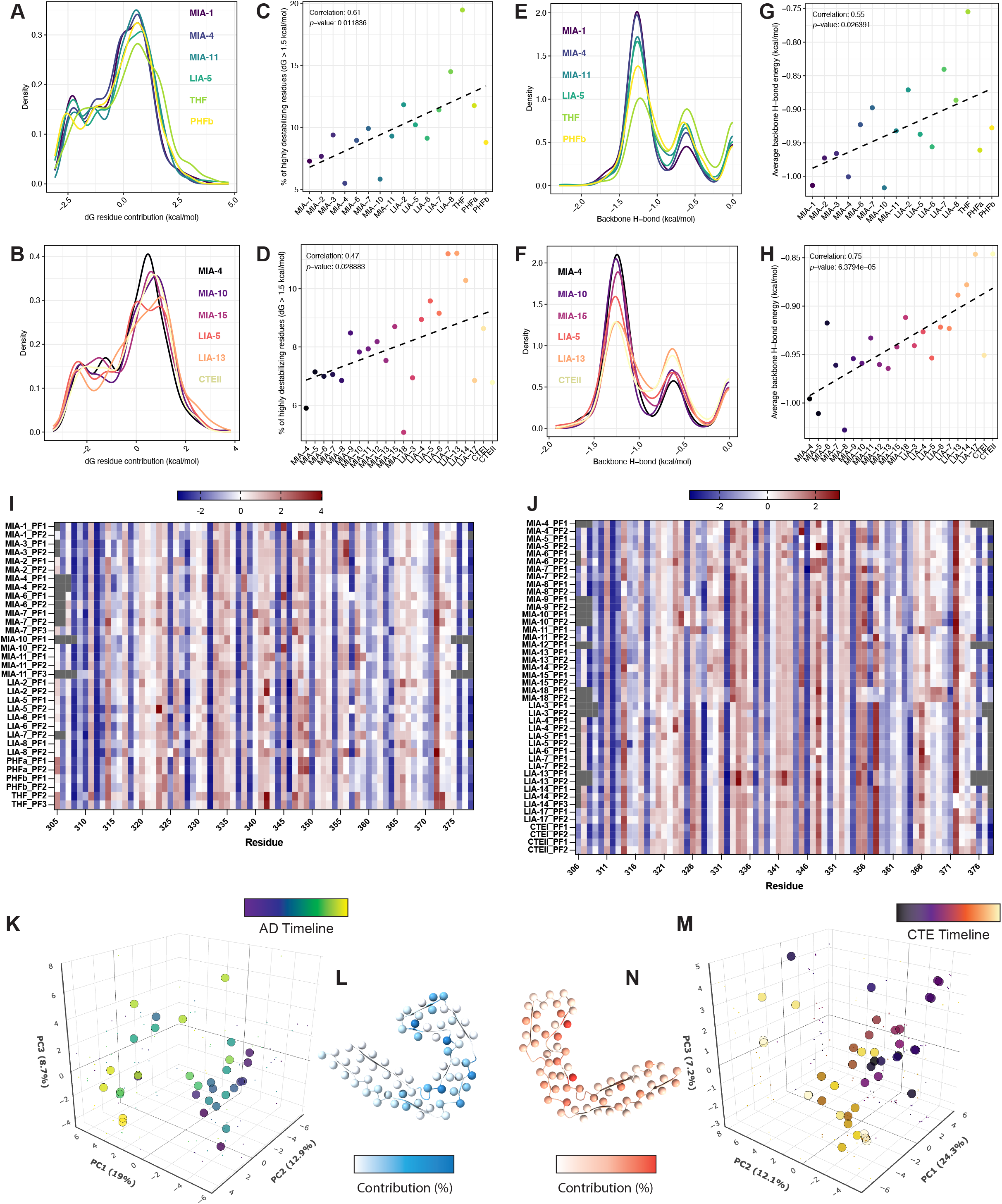
Thermodynamic profiling of recombinant tau intermediate protofilaments. (A, B) Density plots of residue-wise folding energy contributions (ΔG in kcal/mol) for selected structures across the AD (A) and CTE (B) trajectories. (C, D) Percentage of highly destabilizing residues (ΔG > 1.5 kcal/mol) across all modeled structures in the AD (C) and CTE (D) trajectories plotted against the maturation order, with each point representing one structure. A significant positive correlation is observed in both AD and CTE, indicating increasing energetic destabilization with structural progression. Dashed line indicates linear regression. Pearson correlation coefficients and P values are indicated. (E, F) Density plots of backbone hydrogen bond energies (kcal/mol) across a representative subset of structures from AD (E) and CTE (F) trajectories. (G, H) Correlation between average backbone H-bond energy and maturation stage in the AD (G) and CTE (H) trajectories. Progressive weakening of backbone H-bonding is observed in both cases. Dashed line indicates linear regression. Pearson correlation coefficients and P values are indicated. (I-J) Thermodynamic profiling of (I) AD and (J) CTE recombinant tau protofilaments. (K-M) Principal component analysis recapitulates the timeline of protofilaments formed in conditions that reproduce the (K-L) AD and (M-N) CTE fold. In plots A-H, K and M, timepoints are color-coded according to the AD (Figure S2 A) and CTE (Figure S2 C) maturation trajectories.

Energetic maps of intermediate (J-shaped) and mature (C-shaped) protofilaments show that stabilizing energy remains concentrated in motifs such as _306_VQIVYK_311_ and _350_PAM4_362_, which are well-known APRs implicated in tau aggregation_46_ (Fig. 3I, J). In contrast, structural frustration accumulates in neighboring sequences, particularly in flexible turn-forming motifs like the PGGG repeats. Importantly, dimensionality reduction analysis of the maps using PCA captures the chronological progression of structural maturation for both AD (Fig. 3K) and CTE fibril trajectories (Fig. 3M). Variance decomposition, which allows identifying the most influential residues, further highlights key stabilizing residues within APRs and destabilizing ones in adjacent PGGG-containing turns as primary contributors to energetic divergence during maturation (Fig. 3L, N).

Taking together, these results suggest that tau maturation, like IAPP, is anchored by APRs. However, unlike IAPP, it is accompanied by a buildup of internal strain in regions flanking these stabilizing cores. Given that fibril formation must ultimately proceed downhill in free energy, this paradox implies the presence of missing stabilizing contributions that the current energetic model fails to capture and would potentially offset the accumulating strain, allowing these polymorphs to fully mature.

### AD and CTE strains emerge from a common pathway via local energetic reorganization

To investigate how polymorphic diversity arises from tau’s maturation pathway, we performed PCA on the residue-level energy profiles of both J-shaped and C-shaped protofilaments. This analysis revealed three distinct clusters: a shared set of early intermediates, and two separate groups corresponding to the mature AD and CTE folds (Fig. 4A). These findings suggest that both tau strains follow a common initial maturation trajectory before diverging into distinct polymorphs late in assembly. Energetic differences between the mature AD and CTE structures are concentrated in the 343–358 region (Fig. 4B). For example, K343 lies in a loop that is considerably tighter in the CTE fold, forcing its side chain into a more destabilizing orientation than in the AD fold. Conversely, R349 adopts a more constrained conformation in the AD fold, generating an unfavorable local energy that is not present in CTE. The most prominent difference centers on an intra-sheet interaction unique to the AD fold: K353 forms a stabilizing salt bridge with D358, supported by exposure of S356 to the solvent. In contrast, this interaction is absent in CTE protofilaments, where D358 remains unsatisfied and S356 is buried within the fibril core^47^. These subtle shifts redistribute local strain, contributing to fold-specific stabilization patterns (Fig. 3I, J).

**Figure 4.**
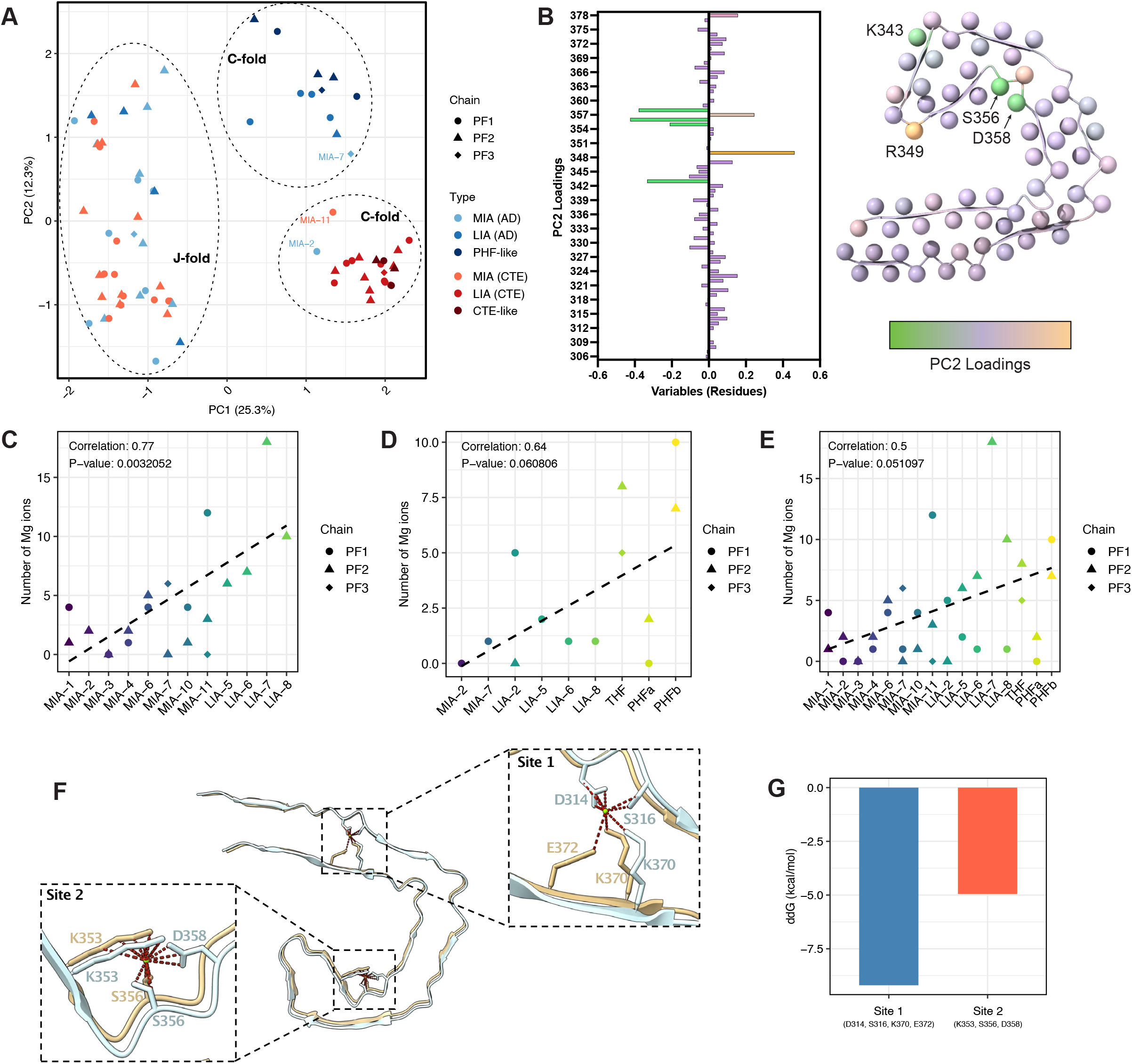
Defining features of the AD and CTE maturation pathways. (A) Principal component analysis of their energetic profiles reveals separate clusters for AD-shaped and CTE-shaped protofilaments and validates a common cluster of intermediate J-folds. Points represent individual protofilaments (PF1, PF2, or PF3) and are colored by structural type. Dashed lines highlight hierarchical clustering of the individual energy profiles, which was performed on the embedding coordinates, and the dendrogram was cut to define three clusters. Labeled protofilaments represent outliers that were misclassified relative to their structural type. (B) Mapping of the second principal component loadings that separate the AD and CTE clusters reveals key residues. (C-E) Prediction of Mg ion binding for fibrils of the AD timeline. Maturation of (C) J-fold, (D) C-fold, and (E) all protofilament structures together, shows a strong correlation of increased metal binding as kinetics progress. Points represent individual protofilaments (PF1, PF2, or PF3), while dashed line indicates linear regression. Pearson correlation coefficients and P values are indicated. Timepoints are color-coded according to the AD maturation trajectory (Figure S2 A). (F) Graphical representation of key metal binding sites in the tau AD C-fold. (G) Metal binding significantly improves the local energetics of the binding sites.

Importantly, these features begin to stratify early-stage intermediates by lineage. In particular, the exposure or burial of S356 appears to act as an early thermodynamic bifurcation point: S356 is mostly exposed in AD-like J-shaped protofilaments, whereas it is largely buried in their CTE-like counterparts (Fig. S5). This suggests that the divergence of AD- and CTE-like fibrils arises gradually from a common maturation pathway, driven by localized energetic reorganization rather than a discrete conformational switch.

### Cofactors redistribute and compensate for protofilament energetic frustration

Because tau fibril maturation is thermodynamically constrained, the observed buildup of internal frustration in both the AD and CTE fold, especially in late-stage structures, implies the presence of stabilizing contributions not captured by protein-only models. Notably, the AD and CTE fibrils analyzed here were assembled under distinct buffer conditions: AD fibrils in the presence of Mg^2+^, and CTE fibrils with NaCl^37,45^. This raised the possibility that extrinsic cofactors such as metal ions help stabilize energetically strained conformations.

To explore this, we predicted metal-binding sites across the tau maturation series^48^. In AD fibrils, Mg^2+^ coordination becomes increasingly favorable as maturation proceeds (Fig. 4C– E). Two key binding sites emerge: one bridging the N- and C-terminal regions of the core, and the other centers on residues S356 and E358 (Fig. 4F). Notably, the latter site only emerges in the final C-shaped AD protofilament and involves S356, the same residue that differentiates the AD and CTE folds (Fig. 4B). This suggests that Mg^2+^ may either stabilize a conformation that emerges with increasing strain, or alternatively, guide the maturation pathway toward that conformation by redistributing energetic strain as it accumulates. Consistent with this model, energetic re-evaluation of the AD mature fibrils with Mg^2+^ ions modeled into these predicted sites confirms that local backbone strain is reduced, and overall energetic profiles improve (Fig. 4G). In contrast, our analysis did not predict Na^+^ binding sites in the CTE-like fibrils. This may reflect both the monovalent and diffuse nature of Na^+^ ions, which are less likely to form stable, geometry-specific coordination complexes. In comparison, Mg^2+^, as a bivalent cation, has a stronger and more predictable binding profile, enabling more confident identification of stabilizing interactions. These differences suggest that, under the present in vitro conditions, AD stabilization likely involves direct metal coordination, whereas CTE stabilization may rely more on nonspecific ionic screening.

To further examine how external cofactors influence fibril energetics, we analyzed a recent structural time series capturing the remodeling of α-synuclein fibrils upon heparin binding^38^ (Fig. S6). Heparin is frequently used in vitro to accelerate α-synuclein fibril formation and promote the emergence of defined polymorphs. Thus, this dataset provides a parallel opportunity for investigating how polyanions influence energetic remodeling during assembly. Early heparin-bound polymorphs (Hep-remod-1 and Hep-remod-2) show enhanced β-structure and improved energetic profiles compared to the apo state (Fig. 5A). However, prolonged heparin exposure leads to the formation of a more frustrated polymorph (Hep-remod-3), as evidenced by a drop in hydrogen-bond stability and a decrease in the fraction of stabilizing residues (Fig. 5B, C). PCA of energy profiles separates early and late heparin-bound states (Fig. 5D), with energetic shifts mapping to known heparin-binding regions (residues 38–45 and 58– 67) (Fig. 5E, F). Since our computational framework cannot directly model polyanions like heparin, the apparent destabilization of the late heparin-induced polymorph likely reflects missing stabilizing contacts that heparin would normally provide.

**Figure 5.**
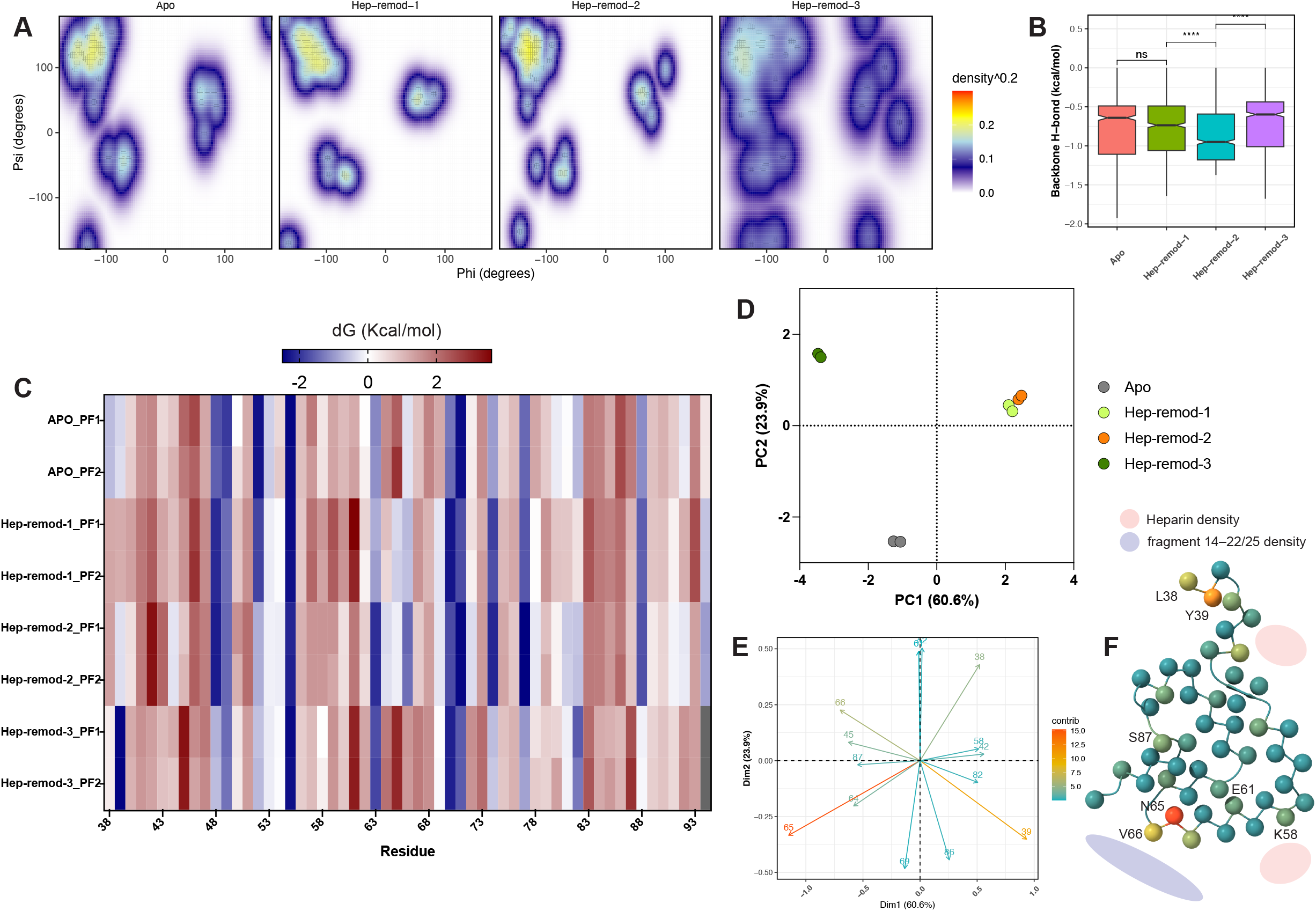
Analysis of heparin-induced aS protofibril restructuring. (A) Ramachandran density plots of aS fibril structures. (B) Backbone hydrogen bond comparison of aS fibril structures. Statistics: t-test comparison of means. (C) Thermodynamic profiling of aS protofilaments. (D) Principal component analysis of aS thermodynamic profiles. (E) Variables plot and (F) mapping of loadings on the aS protofilament core.

Together, these observations show that amyloid fibril maturation, while ultimately thermodynamically downhill, can involve increasing internal frustration that must be offset by stabilizing cofactors. Metal ions and polyanions act as external energetic supports, allowing strained polymorphs to persist by redistributing free energy across the fibril structure.

## Discussion

Amyloid fibrils are defined by their shared cross-β architecture but exhibit remarkable structural diversity, giving rise to polymorphs with distinct biological and pathological properties. Despite the accumulation of many high-resolution cryo-EM structures, the thermodynamic principles that govern how specific polymorphs emerge, evolve, and stabilize over time remain poorly understood. This knowledge gap is especially pronounced in the context of fibril maturation, where assembly occurs over time and is shaped not only by sequence-encoded features but also by environmental cofactors that are often overlooked in structural models.

Here, we address this gap by combining energetic profiling^6,33^ with time-resolved cryo-EM datasets from three amyloid-forming proteins: IAPP, tau, and α-synuclein. Across more than 110 protofilament structures, we uncover a set of unifying thermodynamic features that shape fibril assembly, and we identify system-specific deviations that reveal how intrinsic and extrinsic factors interact to produce diverse polymorphic outcomes.

A central finding is the consistent role of APRs^49^ as thermodynamic anchors during amyloid maturation^5,50-55^. In all systems studied, these short segments dominate the stabilizing energy landscape and expand their structural influence as fibrils mature. This pattern is particularly striking in IAPP, where progressive stabilization of three APRs drives divergence into distinct structural polymorphs that, despite their morphological differences, achieve equivalent overall thermodynamic stability. This convergence echoes classical structure–function studies showing that targeted substitutions at L12,⍰F15,⍰V17, and⍰I26 markedly reduce β-sheet content, suppress fibril growth, and alleviate β-cell cytotoxicity in vitro and in rodent models^39,42,43,56-58^. Comparative genomics strengthens the point: rodents, whose IAPP is naturally non-amyloidogenic, harbor exactly such protective substitutions⍰^59^, and engineering the same changes into human IAPP converts it into a potent, non-aggregating dominant-negative inhibitor of the wild-type peptide^60,61^. These insights have already been translated into clinic-approved analogues that retain glycemic control yet resists fibrillation^62^. Our findings support a model in which APRs act as reusable energetic cores, adaptable across folds and conformational contexts, and essential for nucleating and sustaining ordered amyloid assembly.

In contrast, tau fibrils follow a distinct trajectory. While APRs remain key stabilizing features, the overall maturation process is accompanied by increasing backbone strain and growing energetic frustration, especially in flexible regions such as PGGG-containing turns. This creates a thermodynamic paradox: if amyloid maturation must proceed downhill in free energy, how can it culminate in more frustrated structures? The answer, we propose, lies in stabilizing interactions not accounted for in a protein-only model. This interpretation is supported by our analysis of cofactor interactions. In tau fibrils assembled under Mg^2+^ conditions, predicted metal-binding sites are located in regions already stabilized by APRs, including a segment shown to impart stability specifically to AD-folded tau filaments^63^, and implicated in promoting the prion-like propensity of C-shaped tau strains^64^ and tau polymorph divergence^46^. While these regions are not themselves highly frustrated, metal coordination may modulate the energetic balance of adjacent frustrated segments, helping the structure accommodate strain without global destabilization. Conversely, tau fibrils formed in NaCl do not exhibit structured ion binding, consistent with a reliance on nonspecific ionic screening or other unmodeled interactions.

Interestingly, we identified K343 and R349 as key residues that stratify tau intermediates across maturation and the AD versus CTE timelines. Both residues are consistently destabilized in the two end-state folds. However, they reside in regions where undefined cryo-EM densities have been observed in structures derived from patient tissue, pointing to the possible involvement of unmodeled cofactors or modifications. These sites have the potential to be further stabilized in vivo through post-translational modifications^31^ and lipid-based anionic interactions^65^, which may help alleviate local frustration and contribute to polymorphic divergence. Similarly, S356, a residue that already distinguishes early J-shaped protofilaments by lineage, is often found phosphorylated and serves as a biomarker of pre-tangle soluble tau assemblies in AD^63,66,67^.

In the same light, we found that heparin restructures local hydrogen bonding and alters the fractions of residues adopting stabilizing versus frustrated conformations in aS fibrils, with strain-classifying thermodynamic changes observed in previously identified APRs that mediate aS self-assembly^68^. Whether cofactors stabilize strained structures after they emerge or help guide folding toward them during assembly remains an open question. However, our results clearly show that cofactors shift the distribution of free energy across the fibril and help determine which conformational states are accessible under specific conditions.

We propose a general thermodynamic framework in which sequence-encoded APRs define the energetic backbone of fibril structures, while environmental factors modulate local frustration to favor one polymorph over another. This model explains how structurally distinct polymorphs can emerge from a single sequence, why in vitro and in vivo fibrils often differ, and how disease-specific strain properties may arise from context-dependent stabilization. It also offers a conceptual basis for understanding how small environmental changes, such as ionic conditions, cofactors, and polyanions, can exert outsized effects on amyloid structure, propagation, and toxicity. Methodologically, this work demonstrates the power of combining structural time series with energetic modeling to reconstruct the complex maturation timelines of diverse amyloid systems. The ability to map stabilizing and frustrating residues along assembly trajectories provides a powerful tool for pinpointing residues that act as conformational switches or sites of cofactor dependence. These residues represent promising targets for therapeutic strategies aimed at controlling strain emergence through mutation, ligand binding, or cofactor modulation.

In conclusion, this study provides a thermodynamic blueprint for amyloid polymorphism: a model in which structure emerges from the dynamic balance of intrinsic stability and environmental influence, offering a unified framework to explain polymorphic diversity and its relevance to disease.

## Materials & Methods

### Collection of amyloid fibril structures

We collected all available cryo-EM structures of amyloid fibrils from, to the best of our knowledge, the only three published studies that investigate amyloid formation, maturation and remodeling using a time-course approach. These include: (i) the in vitro maturation of IAPP-S20G over time^36^, (ii) the in vitro assembly of tau into paired helical filaments (PHFs) and chronic traumatic encephalopathy-like filaments^37^, and (iii) the time-dependent remodeling of mature α-synuclein fibrils upon heparin binding^38^.

### Energy profiling of amyloid structures

To assess the thermodynamic stability of amyloid fibrils, we used the FoldX force field^69^. FoldX estimates the free energy of a protein structure by calculating the contribution of each atom based on its interactions with neighboring atoms. These atomic contributions are first summed at the residue level and later at the level of the entire protein. This allows for mapping the per-residue contribution to the total free energy (called ΔG_contrib_), along with detailed breakdowns of individual energetic components. These include van der Waals interactions (ΔG_vdw_), solvation energies for polar and apolar groups (ΔG_solvP_ and ΔG_solvH_), electrostatic interactions (ΔG_el_), hydrogen bonding (ΔG_Hbond_), entropic penalties for main and side chains (ΔS_mc_ and ΔS_sc_), and water-mediated hydrogen bonding (ΔG_wb_).

Since cryo-EM-derived fibril structures vary in the number of stacked monomer layers, all structures were extended to a uniform depth of 10 monomers prior to analysis. Structures were then subjected to side-chain energy minimization using the FoldX RepairPDB command. This procedure optimizes side-chain conformations by sampling from a rotamer library derived from high-resolution X-ray structures, selecting the lowest-energy rotamer for each residue. Subsequently, we used the SequenceDetail function in FoldX to compute per-residue free energy contributions (ΔG_contrib_) and the associated thermodynamic components. To avoid boundary artifacts caused by missing head-to-tail stacking interactions at the fibril termini, monomers located at both ends of the extended fibril were excluded from downstream analysis.

### Removal of low-quality amyloid fibrils structures

The accuracy of the free energy estimation by FoldX correlates with structure quality parameters, such as resolution. Therefore, we defined an empirical partitioning boundary based on a linear relationship between structural resolution and van der Waals (VDW) clashes (Fig. S2). Specifically, structures were excluded if they fell above the line defined by the equation:

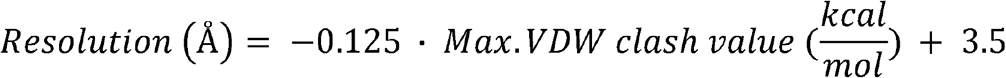

This criterion effectively removes models with both poor resolution and high steric clashes, retaining only high-quality structures for downstream analysis. Resolution values were obtained from the original studies using the Fourier Shell Correlation (FSC) 0.143 criterion^70^, which estimates the resolution of the reconstructed cryo-EM maps. Van der Waals clashes were identified using FoldX.

As a result of this filtering strategy, the following models were excluded from the analysis: MIA-5, MIA-8, and LIA-4 (from the in vitro assembly of tau into paired helical filaments), and MIA-1, MIA-3, and MIA-14 (from the in vitro assembly of tau into CTE-like filaments). Additionally, the first protofilament of LIA-7 and of THF were removed due to excessive steric clashes, while the remaining protofilaments in these structures were retained for analysis.

### Dimensionality reduction analysis

To investigate which residues are energetically driving the maturation and remodeling of amyloid fibrils over time, we performed principal component analysis (PCA) on the per-residue free energy contributions (ΔG_contrib_) calculated by FoldX. These values were compiled into a matrix where rows represented individual protofilaments and columns corresponded to aligned residue positions. Prior to dimensionality reduction, the data was normalized by row-wise standardization to account for differences in absolute energy scales across protofilaments. Then, missing values, which occurred in some protofilaments due to unresolved residues, were imputed using the imputePCA function from the missMDA R package with regularized iterative PCA^71^. PCA was then performed on the imputed, standardized matrix using the prcomp function from the stats R package. Clustering was subsequently applied to the first three principal components using hierarchical clustering via the hclust function.

### Metal binding analysis

To investigate the potential contribution of metal ions to the in vitro assembly of tau fibrils, we used a previously established high-accuracy algorithm implemented in FoldX^48^ to predict metal binding sites in each protofilament. This method identifies positions on the protein surface where metal ions can form favorable electrostatic interactions with multiple protein atoms in geometrically optimal configurations. This approach has been shown to predict more than 90% of the metal binding sites in globular proteins with a positional accuracy of ≤0.6 Å^48^. Given that the tau (297–391) construct forms paired helical filaments (PHFs) when incubated in phosphate buffer supplemented with magnesium chloride^37,45^, we predicted magnesium binding sites in all protofilaments derived from the PHF reaction. In contrast, for structures derived from the CTE-like reaction, which uses sodium chloride instead of magnesium chloride^37,45^, we predicted sodium binding sites (although no binding sites with high affinity could be inferred).

To assess the impact of predicted metal binding on structural energetics, we incorporated the identified metal ions into each structure and recalculated per-residue free energy contributions (ΔG_contrib_) using the FoldX SequenceDetail function.

### Quantification, statistics and visualizations

The methods of statistical analysis are provided in detail in the corresponding figure legends. Visualizations were performed with GraphPad prism or custom R scripts using the packages ggplot2 and plotly. ChimeraX was used to visualize protein structures^72^.

## Data Availability

All data generated in this study are available as Source data and are also available on the Zenodo database under accession code XXXXX.

## References

1. Chiti, F. & Dobson, C.M. Protein Misfolding, Amyloid Formation, and Human Disease: A Summary of Progress Over the Last Decade. Annu Rev Biochem 86, 27–68 (2017). 10.1146/annurev-biochem-061516-045115

2. Buxbaum, J. N. et al. Amyloid nomenclature 2024: update, novel proteins, and recommendations by the International Society of Amyloidosis (ISA) Nomenclature Committee. Amyloid 31, 249–256 (2024). 10.1080/13506129.2024.2405948

3. Nelson, R. et al. Structure of the cross-beta spine of amyloid-like fibrils. Nature 435, 773-778 (2005). 10.1038/nature03680

4. Sawaya, M. R., Hughes, M. P., Rodriguez, J. A., Riek, R. & Eisenberg, D. S. The expanding amyloid family: Structure, stability, function, and pathogenesis. Cell 184, 4857–4873 (2021). 10.1016/j.cell.2021.08.013

5. Louros, N., Schymkowitz, J. & Rousseau, F. Mechanisms and pathology of protein misfolding and aggregation. Nature Reviews Molecular Cell Biology 24, 912–933 (2023).

6. van der Kant, R., Louros, N., Schymkowitz, J. & Rousseau, F. Thermodynamic analysis of amyloid fibril structures reveals a common framework for stability in amyloid polymorphs. Structure 30, 1178-1189.e1173 (2022). 10.1016/j.str.2022.05.002

7. Fernandez-Escamilla, A. M., Rousseau, F., Schymkowitz, J. & Serrano, L. Prediction of sequence-dependent and mutational effects on the aggregation of peptides and proteins. Nat Biotechnol 22, 1302–1306 (2004). 10.1038/nbt1012

8. Rousseau, F., Serrano, L. & Schymkowitz, J. W. How evolutionary pressure against protein aggregation shaped chaperone specificity. J Mol Biol 355, 1037–1047 (2006). 10.1016/j.jmb.2005.11.035

9. Krammer, C., Schätzl, H. M. & Vorberg, I. Prion-like propagation of cytosolic protein aggregates: insights from cell culture models. Prion 3, 206–212 (2009). 10.4161/pri.3.4.10013

10. Goedert, M., Clavaguera, F. & Tolnay, M. The propagation of prion-like protein inclusions in neurodegenerative diseases. Trends in Neurosciences 33, 317–325 (2010). 10.1016/j.tins.2010.04.003

11. Sanders, D. W. et al. Distinct tau prion strains propagate in cells and mice and define different tauopathies. Neuron 82, 1271–1288 (2014). 10.1016/j.neuron.2014.04.047

12. Goedert, M. Alzheimer’s and Parkinson’s diseases: The prion concept in relation to assembled Aβ, tau, and α-synuclein. Science 349, 1255555 (2015). 10.1126/science.1255555

13. Willbold, D., Strodel, B., Schröder, G. F., Hoyer, W. & Heise, H. Amyloid-type Protein Aggregation and Prion-like Properties of Amyloids. Chemical Reviews 121, 8285–8307 (2021). 10.1021/acs.chemrev.1c00196

14. Shahnawaz, M. et al. Discriminating α-synuclein strains in Parkinson’s disease and multiple system atrophy. Nature 578, 273–277 (2020). 10.1038/s41586-020-1984-7

15. Gallardo, R., Ranson, N. A. & Radford, S. E. Amyloid structures: much more than just a cross-β fold. Curr Opin Struct Biol 60, 7–16 (2020). 10.1016/j.sbi.2019.09.001

16. Louros, N., Schymkowitz, J. & Rousseau, F. Heterotypic amyloid interactions: Clues to polymorphic bias and selective cellular vulnerability? Current Opinion in Structural Biology 72, 176–186 (2022). 10.1016/j.sbi.2021.11.007

17. Scheres, S. H. W., Ryskeldi-Falcon, B. & Goedert, M. Molecular pathology of neurodegenerative diseases by cryo-EM of amyloids. Nature 621, 701–710 (2023). 10.1038/s41586-023-06437-2

18. Shi, Y. et al. Structure-based classification of tauopathies. Nature 598, 359–363 (2021). 10.1038/s41586-021-03911-7

19. Yang, Y. et al. Cryo-EM structures of amyloid-β 42 filaments from human brains. Science 375, 167–172 (2022). 10.1126/science.abm7285

20. Kollmer, M. et al. Cryo-EM structure and polymorphism of Aβ amyloid fibrils purified from Alzheimer’s brain tissue. Nat Commun 10, 4760 (2019). 10.1038/s41467-019-12683-8

21. Yang, Y. et al. Cryo-EM structures of Aβ40 filaments from the leptomeninges of individuals with Alzheimer’s disease and cerebral amyloid angiopathy. Acta Neuropathologica Communications 11, 191 (2023). 10.1186/s40478-023-01694-8

22. Schweighauser, M. et al. Structures of α-synuclein filaments from multiple system atrophy. Nature 585, 464–469 (2020). 10.1038/s41586-020-2317-6

23. Yang, Y. et al. Structures of α-synuclein filaments from human brains with Lewy pathology. Nature 610, 791–795 (2022). 10.1038/s41586-022-05319-3

24. Arseni, D. et al. TDP-43 forms amyloid filaments with a distinct fold in type A FTLD-TDP. Nature 620, 898–903 (2023). 10.1038/s41586-023-06405-w

25. Arseni, D. et al. Heteromeric amyloid filaments of ANXA11 and TDP-43 in FTLD-TDP type C. Nature 634, 662–668 (2024). 10.1038/s41586-024-08024-5

26. Schmidt, M. et al. Cryo-EM structure of a transthyretin-derived amyloid fibril from a patient with hereditary ATTR amyloidosis. Nature Communications 10, 5008 (2019). 10.1038/s41467-019-13038-z

27. Radamaker, L. et al. Cryo-EM structure of a light chain-derived amyloid fibril from a patient with systemic AL amyloidosis. Nature Communications 10, 1103 (2019). 10.1038/s41467-019-09032-0

28. Lövestam, S. et al. Seeded assembly in vitro does not replicate the structures of α-synuclein filaments from multiple system atrophy. FEBS Open Bio 11, 999–1013 (2021). 10.1002/2211-5463.13110

29. Zhang, W. et al. Heparin-induced tau filaments are polymorphic and differ from those in Alzheimer’s and Pick’s diseases. Elife 8 (2019). 10.7554/eLife.43584

30. Konstantoulea, K. et al. Heterotypic Amyloid β interactions facilitate amyloid assembly and modify amyloid structure. Embo j 41, e108591 (2022). 10.15252/embj.2021108591

31. Arakhamia, T. et al. Posttranslational Modifications Mediate the Structural Diversity of Tauopathy Strains. Cell 180, 633-644.e612 (2020). 10.1016/j.cell.2020.01.027

32. Mullapudi, V. et al. Network of hotspot interactions cluster tau amyloid folds. Nature Communications 14, 895 (2023). 10.1038/s41467-023-36572-3

33. Louros, N., van der Kant, R., Schymkowitz, J. & Rousseau, F. StAmP-DB: a platform for structures of polymorphic amyloid fibril cores. Bioinformatics 38, 2636–2638 (2022). 10.1093/bioinformatics/btac126

34. Mahmoudinobar, F., Urban, J. M., Su, Z., Nilsson, B. L. & Dias, C. L. Thermodynamic Stability of Polar and Nonpolar Amyloid Fibrils. Journal of Chemical Theory and Computation 15, 3868–3874 (2019). 10.1021/acs.jctc.9b00145

35. Larsen, J. A. et al. The mechanism of amyloid fibril growth from Φ-value analysis. Nat Chem 17, 403–411 (2025). 10.1038/s41557-024-01712-9

36. Wilkinson, M. et al. Structural evolution of fibril polymorphs during amyloid assembly. Cell 186, 5798-5811.e5726 (2023). 10.1016/j.cell.2023.11.025

37. Lövestam, S. et al. Disease-specific tau filaments assemble via polymorphic intermediates. Nature 625, 119–125 (2024). 10.1038/s41586-023-06788-w

38. Tao, Y. et al. Time-course remodeling and pathology intervention of α-synuclein amyloid fibril by heparin and heparin-like oligosaccharides. Nature Structural & Molecular Biology 32, 369–380 (2025). 10.1038/s41594-024-01407-2

39. Louros, N. N. et al. Structural studies and cytotoxicity assays of “aggregation-prone” IAPP(8-16) and its non-amyloidogenic variants suggest its important role in fibrillogenesis and cytotoxicity of human amylin. Biopolymers 104, 196–205 (2015). 10.1002/bip.22650

40. Mirecka, E. A. et al. β-Hairpin of Islet Amyloid Polypeptide Bound to an Aggregation Inhibitor. Scientific Reports 6, 33474 (2016). 10.1038/srep33474

41. Westermark, P., Engström, U., Johnson, K. H., Westermark, G. T. & Betsholtz, C. Islet amyloid polypeptide: pinpointing amino acid residues linked to amyloid fibril formation. Proceedings of the National Academy of Sciences 87, 5036–5040 (1990). 10.1073/pnas.87.13.5036

42. Abedini, A. & Raleigh, D. P. Destabilization of Human IAPP Amyloid Fibrils by Proline Mutations Outside of the Putative Amyloidogenic Domain: Is There a Critical Amyloidogenic Domain in Human IAPP? Journal of Molecular Biology 355, 274–281 (2006). 10.1016/j.jmb.2005.10.052

43. Fox, A. et al. Selection for nonamyloidogenic mutants of islet amyloid polypeptide (IAPP) identifies an extended region for amyloidogenicity. Biochemistry 49, 7783–7789 (2010). 10.1021/bi100337p

44. Nilsson, M. R. & Raleigh, D. P. Analysis of amylin cleavage products provides new insights into the amyloidogenic region of human amylin. J Mol Biol 294, 1375–1385 (1999). 10.1006/jmbi.1999.3286

45. Lövestam, S. et al. Assembly of recombinant tau into filaments identical to those of Alzheimer’s disease and chronic traumatic encephalopathy. eLife 11, e76494 (2022). 10.7554/eLife.76494

46. Louros, N. et al. Local structural preferences in shaping tau amyloid polymorphism. Nature Communications 15, 1028 (2024). 10.1038/s41467-024-45429-2

47. Goedert, M. & Spillantini, M. G. Ordered Assembly of Tau Protein and Neurodegeneration. Adv Exp Med Biol 1184, 3–21 (2019). 10.1007/978-981-32-9358-8_1

48. Schymkowitz, J. W. et al. Prediction of water and metal binding sites and their affinities by using the Fold-X force field. Proc Natl Acad Sci U S A 102, 10147–10152 (2005). 10.1073/pnas.0501980102

49. Langenberg, T. et al. Thermodynamic and Evolutionary Coupling between the Native and Amyloid State of Globular Proteins. Cell Reports 31 (2020). 10.1016/j.celrep.2020.03.076

50. Teng, P. K. & Eisenberg, D. Short protein segments can drive a non-fibrillizing protein into the amyloid state. Protein Eng Des Sel 22, 531–536 (2009). 10.1093/protein/gzp037

51. Ventura, S. et al. Short amino acid stretches can mediate amyloid formation in globular proteins: The Src homology 3 (SH3) case. Proceedings of the National Academy of Sciences 101, 7258–7263 (2004). 10.1073/pnas.0308249101

52. Louros, N. et al. WALTZ-DB 2.0: an updated database containing structural information of experimentally determined amyloid-forming peptides. Nucleic Acids Res 48, D389–D393 (2020). 10.1093/nar/gkz758

53. Louros, N., Orlando, G., De Vleeschouwer, M., Rousseau, F. & Schymkowitz, J. Structure-based machine-guided mapping of amyloid sequence space reveals uncharted sequence clusters with higher solubilities. Nature communications 11, 3314 (2020). 10.1038/s41467-020-17207-3

54. Sawaya, M. R. et al. Atomic structures of amyloid cross-β spines reveal varied steric zippers. Nature 447, 453–457 (2007). 10.1038/nature05695

55. Wagner, J. et al. Medin co-aggregates with vascular amyloid-beta in Alzheimer’s disease. Nature 612, 123–131 (2022). 10.1038/s41586-022-05440-3

56. Bernhardt, N. A., Berhanu, W. M. & Hansmann, U. H. Mutations and seeding of amylin fibril-like oligomers. J Phys Chem B 117, 16076–16085 (2013). 10.1021/jp409777p

57. Abedini, A., Meng, F. & Raleigh, D. P. A Single-Point Mutation Converts the Highly Amyloidogenic Human Islet Amyloid Polypeptide into a Potent Fibrillization Inhibitor. Journal of the American Chemical Society 129, 11300–11301 (2007). 10.1021/ja072157y

58. Louros, N. N. et al. Tracking the amyloidogenic core of IAPP amyloid fibrils: Insights from micro-Raman spectroscopy. J Struct Biol 199, 140–152 (2017). 10.1016/j.jsb.2017.06.002

59. Betsholtz, C. et al. Sequence divergence in a specific region of islet amyloid polypeptide (IAPP) explains differences in islet amyloid formation between species. FEBS Letters 251, 261–264 (1989). 10.1016/0014-5793(89)81467-X

60. Manchanda, A. & Goyal, B. Deciphering the impact of F23L mutation on the aggregation propensity of human islet amyloid polypeptide using molecular simulations. Journal of Molecular Liquids 411, 125775 (2024). 10.1016/j.molliq.2024.125775

61. Abedini, A., Meng, F. & Raleigh, D. P. A single-point mutation converts the highly amyloidogenic human islet amyloid polypeptide into a potent fibrillization inhibitor. J Am Chem Soc 129, 11300–11301 (2007). 10.1021/ja072157y

62. Ridgway, Z. et al. Analysis of Proline Substitutions Reveals the Plasticity and Sequence Sensitivity of Human IAPP Amyloidogenicity and Toxicity. Biochemistry 59, 742–754 (2020). 10.1021/acs.biochem.9b01109

63. Leonard, C., Phillips, C. & McCarty, J. Insight Into Seeded Tau Fibril Growth From Molecular Dynamics Simulation of the Alzheimer’s Disease Protofibril Core. Front Mol Biosci 8, 624302 (2021). 10.3389/fmolb.2021.624302

64. Fitzpatrick, A. W. P. et al. Cryo-EM structures of tau filaments from Alzheimer’s disease. Nature 547, 185–190 (2017). 10.1038/nature23002

65. Fowler, S. L. et al. Tau filaments are tethered within brain extracellular vesicles in Alzheimer’s disease. Nature Neuroscience 28, 40–48 (2025). 10.1038/s41593-024-01801-5

66. Islam, T. et al. Phospho-tau serine-262 and serine-356 as biomarkers of pre-tangle soluble tau assemblies in Alzheimer’s disease. Nat Med 31, 574–588 (2025). 10.1038/s41591-024-03400-0

67. Taylor, L. W. et al. p-tau Ser356 is associated with Alzheimer’s disease pathology and is lowered in brain slice cultures using the NUAK inhibitor WZ4003. Acta Neuropathol 147, 7 (2024). 10.1007/s00401-023-02667-w

68. Doherty, C. P. A. et al. A short motif in the N-terminal region of α-synuclein is critical for both aggregation and function. Nat Struct Mol Biol 27, 249–259 (2020). 10.1038/s41594-020-0384-x

69. Schymkowitz, J. et al. The FoldX web server: an online force field. Nucleic Acids Res 33, W382–388 (2005). 10.1093/nar/gki387

70. Rosenthal, P. B. & Henderson, R. Optimal determination of particle orientation, absolute hand, and contrast loss in single-particle electron cryomicroscopy. Journal of molecular biology 333, 721–745 (2003).

71. Josse, J. & Husson, F. missMDA: a package for handling missing values in multivariate data analysis. Journal of statistical software 70, 1–31 (2016).

72. Goddard, T. D. et al. UCSF ChimeraX: Meeting modern challenges in visualization and analysis. Protein Science 27, 14–25 (2018).

