## Supplementary Information for "Energetic profiling reveals thermodynamic principles underlying amyloid fibril maturation"

This PDF file includes:

Supplementary Figures 1-6

Supplementary Table 1

Supplementary References

**Supplementary Figure 1.** Ramachandran density plots of individual protofilaments formed during the structural evolution of IAPP S20G amyloid fibrils^1^.

**Supplementary Figure 2. AD and CTE tau polymorphic intermediate filaments**^2^**.** (A) Overview of fibril structures formed by recombinant truncated tau (297-391) in conditions that reproduce the AD-associated disease paired helical filaments (PHFs). Protofilaments in grey have been removed from the analysis after prefiltering. (B) Prefiltering protofilament structures of the AD timeline. Structures above the line, showing strong residue clash energies (x-axis) or with poor resolution were excluded from further analysis (shown in red). (C) Overview of fibrils formed by recombinant truncated tau (297-391) in conditions that reproduce the CTE-associated disease filaments. (D) Prefiltering protofilament structures of the CTE timeline as in B.

**Supplementary Figure 3. Structural assessment and comparison of tau AD protofilament structures.** (A) Ramachandran density plot analysis of tau AD protofilaments. Densities corresponding to dihedrals in non-beta conformations increase over maturation. (B-E) Density plots indicating the per residue distribution of (B) total, (C) backbone hydrogen bond, (D) and solvation energies, as well as (E) side chain burials. Each curve represents a different structure of the AD timeline (color-coded as in Figure S2 A).

**Supplementary Figure 4. Structural assessment and comparison of tau CTE protofilament structures.** (A) Ramachandran density plot analysis of tau CTE protofilaments. Densities corresponding to dihedrals in non-beta conformations increase over maturation. (B-E) Density plots indicating the per residue distribution of (B) total, (C) backbone hydrogen bond, (D) and solvation energies, as well as (E) side chain burials. Each curve represents a different structure of the CTE timeline (color-coded as in Figure S2 C).

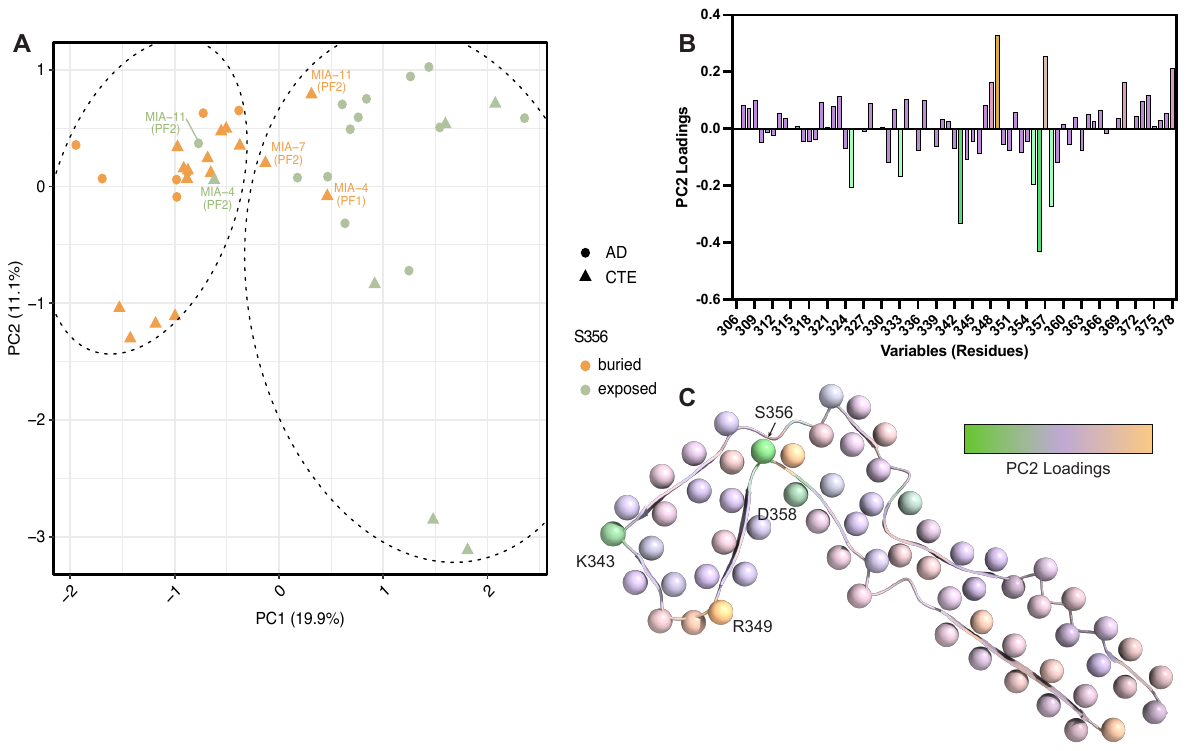
**Supplementary Figure 5. Principal component analysis of energetic profiles of intermediate tau protofilaments.** (A) PCA plot of J-shaped tau protofilaments derived from both the AD (circle-shaped points) and CTE (triangle-shaped points) maturation timelines. The exposure of S356 is a defining feature that clusters all J-shaped protofilaments from both timelines. Dashed lines highlight hierarchical clustering of the individual energy profiles, which was performed on the embedding coordinates, and the dendrogram was cut to define two clusters. Labeled protofilaments represent outliers that were misclassified relative to their structural type. (B) Breakdown of individual loadings of the first principal component. (C) Example of a J-shaped tau protofilament, color-coded based on the individual loadings of the first principal component, reveals that S356, E358 and K343 defined J-shaped protofilaments and are anticorrelated to the energetics of R349 and L357.

**Supplementary Figure 6.** Graphical representation of aS fibril restructuring in the presence of heparin as a co-factor^3^. CryoEM structures were determined at different timepoints following the addition of heparin: before heparin addition (Apo), immediately after mixing (Hep-remod-1, 0h), 1h after mixing (Hep-remod-2, 1h), and after long term mixing (Hep-remod-3, 3 days).

**Supplementary Table 1.** Cryo-EM structures reanalyzed in this study.

| **PDB ID** | **Structure Title** | **Resolution (Å)** | **Structure Author** | **Protein** |
| --- | --- | --- | --- | --- |
| 8Q27 | Tau - AD-MIA1 | 2.02 | Lovestam, S., Scheres, S.H.W., Goedert, M., Li, D. | Tau |
| 8Q2J | Tau - AD-MIA2 | 2.23 | Lovestam, S., Scheres, S.H.W., Goedert, M., Li, D. | Tau |
| 8Q2K | Tau - AD-MIA3 | 2.88 | Lovestam, S., Scheres, S.H.W., Goedert, M., Li, D. | Tau |
| 8Q2L | Tau - AD-MIA4 | 2.20 | Lovestam, S., Scheres, S.H.W., Goedert, M., Li, D. | Tau |
| 8Q7F | Tau - AD-MIA5 | 3.72 | Lovestam, S., Scheres, S.H.W., Goedert, M., Li, D. | Tau |
| 8Q7L | Tau - AD-MIA6 | 2.82 | Lovestam, S., Scheres, S.H.W., Goedert, M., Li, D. | Tau |
| 8Q7M | Tau - AD-MIA7 | 3.26 | Lovestam, S., Scheres, S.H.W., Goedert, M., Li, D. | Tau |
| 8Q7P | Tau - AD-MIA8 | 3.28 | Lovestam, S., Scheres, S.H.W., Goedert, M., Li, D. | Tau |
| 8Q8C | Tau - AD-MIA10 | 1.92 | Lovestam, S., Scheres, S.H.W., Goedert, M., Li, D. | Tau |
| 8Q7T | Tau - AD-MIA11 | 3.00 | Lovestam, S., Scheres, S.H.W., Goedert, M., Li, D. | Tau |
| 8Q88 | Tau - AD-LIA2 | 2.95 | Lovestam, S., Scheres, S.H.W., Goedert, M., Li, D. | Tau |
| 8Q8E | Tau - AD-LIA4 | 3.81 | Lovestam, S., Scheres, S.H.W., Goedert, M., Li, D. | Tau |
| 8Q8F | Tau - AD-LIA5 | 2.93 | Lovestam, S., Scheres, S.H.W., Goedert, M., Li, D. | Tau |
| 8Q8D | Tau - AD-LIA6 | 3.04 | Lovestam, S., Scheres, S.H.W., Goedert, M., Li, D. | Tau |
| 8Q8L | Tau - AD-LIA7 | 3.26 | Lovestam, S., Scheres, S.H.W., Goedert, M., Li, D. | Tau |
| 8QCP | Tau - AD-LIA8 | 3.21 | Lovestam, S., Scheres, S.H.W., Goedert, M., Li, D. | Tau |
| 8Q8S | Tau - AD-THF | 2.68 | Lovestam, S., Scheres, S.H.W., Goedert, M., Li, D. | Tau |
| 8Q8R | Tau - AD-PHFa | 2.10 | Lovestam, S., Scheres, S.H.W., Goedert, M., Li, D. | Tau |
| 8Q8M | Tau - AD-PHFb | 2.95 | Lovestam, S., Scheres, S.H.W., Goedert, M., Li, D. | Tau |
| 8Q8U | Tau - CTE-MIA1 | 3.30 | Lovestam, S., Scheres, S.H.W., Goedert, M., Li, D. | Tau |
| 8Q8V | Tau - CTE-MIA3 | 3.80 | Lovestam, S., Scheres, S.H.W., Goedert, M., Li, D. | Tau |
| 8Q8W | Tau - CTE-MIA4 | 2.85 | Lovestam, S., Scheres, S.H.W., Goedert, M., Li, D. | Tau |
| 8Q8X | Tau - CTE-MIA5 | 2.54 | Lovestam, S., Scheres, S.H.W., Goedert, M., Li, D. | Tau |
| 8Q8Y | Tau - CTE-MIA6 | 2.88 | Lovestam, S., Scheres, S.H.W., Goedert, M., Li, D. | Tau |
| 8Q8Z | Tau - CTE-MIA7 | 3.16 | Lovestam, S., Scheres, S.H.W., Goedert, M., Li, D. | Tau |
| 8Q98 | Tau - CTE-MIA8 | 1.75 | Lovestam, S., Scheres, S.H.W., Goedert, M., Li, D. | Tau |
| 8Q97 | Tau - CTE-MIA9 | 2.99 | Lovestam, S., Scheres, S.H.W., Goedert, M., Li, D. | Tau |
| 8Q99 | Tau - CTE-MIA10 | 2.70 | Lovestam, S., Scheres, S.H.W., Goedert, M., Li, D. | Tau |
| 8Q9A | Tau - CTE-MIA11 | 3.04 | Lovestam, S., Scheres, S.H.W., Goedert, M., Li, D. | Tau |
| 8QCR | Tau - CTE-MIA12 | 2.75 | Lovestam, S., Scheres, S.H.W., Goedert, M., Li, D. | Tau |
| 8Q9B | Tau - CTE-MIA13 | 3.10 | Lovestam, S., Scheres, S.H.W., Goedert, M., Li, D. | Tau |
| 8Q9C | Tau - CTE-MIA14 | 3.40 | Lovestam, S., Scheres, S.H.W., Goedert, M., Li, D. | Tau |
| 8Q9D | Tau - CTE-MIA15 | 3.16 | Lovestam, S., Scheres, S.H.W., Goedert, M., Li, D. | Tau |
| 8Q9E | Tau - CTE-MIA18 | 2.97 | Lovestam, S., Scheres, S.H.W., Goedert, M., Li, D. | Tau |
| 8Q9F | Tau - CTE-LIA3 | 1.91 | Lovestam, S., Scheres, S.H.W., Goedert, M., Li, D. | Tau |
| 8Q9H | Tau - CTE-LIA4 | 2.18 | Lovestam, S., Scheres, S.H.W., Goedert, M., Li, D. | Tau |
| 8Q9G | Tau - CTE-LIA5 | 2.65 | Lovestam, S., Scheres, S.H.W., Goedert, M., Li, D. | Tau |
| 8Q9I | Tau - CTE-LIA6 | 2.56 | Lovestam, S., Scheres, S.H.W., Goedert, M., Li, D. | Tau |
| 8Q9J | Tau - CTE-LIA7 | 2.96 | Lovestam, S., Scheres, S.H.W., Goedert, M., Li, D. | Tau |
| 8Q9K | Tau - CTE-LIA13 | 3.20 | Lovestam, S., Scheres, S.H.W., Goedert, M., Li, D. | Tau |
| 8Q9L | Tau - CTE-LIA14 | 2.76 | Lovestam, S., Scheres, S.H.W., Goedert, M., Li, D. | Tau |
| 8Q9O | Tau - CTE-LIA17 | 3.10 | Lovestam, S., Scheres, S.H.W., Goedert, M., Li, D. | Tau |
| 8Q9M | Tau - CTE type I | 2.65 | Lovestam, S., Scheres, S.H.W., Goedert, M., Li, D. | Tau |
| 8QJJ | Tau - CTE type II | 3.35 | Lovestam, S., Scheres, S.H.W., Goedert, M., Li, D. | Tau |
| 8HZB | Hep-remod-1 | 3.20 | Tao, Y.Q., Zhao, Q.Y., Liu, C., Li, D. | α-synuclein |
| 8HZC | Hep-remod-2 | 3.20 | Tao, Y.Q., Zhao, Q.Y., Liu, C., Li, D. | α-synuclein |
| 8HZS | Hep-remod-3 | 3.30 | Tao, Y.Q., Zhao, Q.Y., Liu, C., Li, D. | α-synuclein |
| 6A6B | alpha-synuclein fiber (apo) | 3.07 | Li, Y.W., Zhao, C.Y., Luo, F., Liu, Z., Gui, X., Luo, Z., Zhang, X., Li, D., Liu, C., Li, X. | α-synuclein |
| 8AWT | IAPP S20G lag-phase fibril polymorph 2PF-P | 3.00 | Wilkinson, M., Xu, Y., Gallardo, R., Radford, S.E., Ranson, N.A. | IAPP |
| 8AZ0 | IAPP S20G growth-phase fibril polymorph 2PF-L | 3.40 | Wilkinson, M., Xu, Y., Gallardo, R., Radford, S.E., Ranson, N.A. | IAPP |
| 8AZ1 | IAPP S20G growth-phase fibril polymorph 2PF-C | 3.10 | Wilkinson, M., Xu, Y., Gallardo, R., Radford, S.E., Ranson, N.A. | IAPP |
| 8AZ2 | IAPP S20G growth-phase fibril polymorph 3PF-CU | 3.40 | Wilkinson, M., Xu, Y., Gallardo, R., Radford, S.E., Ranson, N.A. | IAPP |
| 8AZ3 | IAPP S20G growth-phase fibril polymorph 4PF-CU | 3.40 | Wilkinson, M., Xu, Y., Gallardo, R., Radford, S.E., Ranson, N.A. | IAPP |
| 8AZ4 | IAPP S20G plateau-phase fibril polymorph 2PF-L | 2.20 | Wilkinson, M., Xu, Y., Gallardo, R., Radford, S.E., Ranson, N.A. | IAPP |
| 8AZ5 | IAPP S20G plateau-phase fibril polymorph 4PF-CU | 2.30 | Wilkinson, M., Xu, Y., Gallardo, R., Radford, S.E., Ranson, N.A. | IAPP |
| 8AZ6 | IAPP S20G plateau-phase fibril polymorph 4PF-LU | 3.10 | Wilkinson, M., Xu, Y., Gallardo, R., Radford, S.E., Ranson, N.A. | IAPP |
| 8AZ7 | IAPP S20G plateau-phase fibril polymorph 4PF-LJ | 2.90 | Wilkinson, M., Xu, Y., Gallardo, R., Radford, S.E., Ranson, N.A. | IAPP |

**Supplementary references**

1	Wilkinson, M. *et al.* Structural evolution of fibril polymorphs during amyloid assembly. *Cell* **186**, 5798-5811.e5726 (2023).

2	Lövestam, S. *et al.* Disease-specific tau filaments assemble via polymorphic intermediates. *Nature* **625**, 119-125 (2024).

3	Tao, Y. *et al.* Time-course remodeling and pathology intervention of α-synuclein amyloid fibril by heparin and heparin-like oligosaccharides. *Nature Structural & Molecular Biology* **32**, 369-380 (2025).
